# Contrasting morphological and acoustic trait spaces suggest distinct benefits to participants in mixed-species bird flocks

**DOI:** 10.1101/2024.04.15.589677

**Authors:** A V Abhijith, Samira Agnihotri, Priti Bangal, Anand Krishnan

## Abstract

Mixed-species bird flocks are dynamic associations that exhibit compositional turnover over relatively small timescales. Morphological diversity and foraging behaviour of species in flocks influences the relative benefits and costs of flock participation. In addition, species within flocks are highly acoustically active. However, the role of acoustic signals in flock assembly remains poorly understood. Here, we examined the relationship between acoustic and morphological trait spaces of bird flocks in peninsular India. We found that participant species are generally more similar in body mass than expected by chance. Flocks in general were dominated by smaller-sized species. Conversely, we found that flock participants are not similar in acoustic traits. Much literature suggests that morphology and acoustic signal parameters are known to be correlated, but we present evidence to suggest that these two trait spaces are decoupled at the community scale. This may enable species to derive distinct sets of benefits from both sets of traits, and provides valuable insight into the dynamic processes driving flock assembly.

**Lay summary:** Participants in mixed-species bird flocks tend to group together with similar-sized individuals. This morphological similarity in a crowded flock may result in acoustic signal overlap, as the two sets of traits are correlated to each other. Here, we find evidence to suggest that morphological and acoustic traits are decoupled in these interspecific associations, putatively enabling species to derive benefits from flocking with similar-sized species, and simultaneously minimize acoustic masking interference.

## Introduction

Insectivorous forest birds occupying diverse habitats form interspecific foraging groups that move together. These mixed-species flocks vary in their species composition, size, permanence, and association strength between species pairs (Moynihan 1962, Terborgh 1990, Greenberg 2001, Sridhar *et al*. 2009). Participant bird species in these flocks accrue two key benefits: increased foraging efficiency, and anti-predatory benefits (Sridhar *et al*. 2009). Decades of global studies on flocks have provided varying support for generalizable rules of flock assembly. However, most studies to date focus on understanding the phenotypic composition of flocks, and how this influences the trade-off between benefits and costs from flock participation. For instance, body size of participant species is an important trait that potentially influences the benefits of joining flocks (Sridhar *et al*. 2012, Bangal *et al*. 2021), whereas distinct foraging guilds/patterns (Goodale *et al*. 2020) and differences in foraging behaviour (Sridhar & Shanker 2014) may reduce the cost of competition in flocks. In addition to foraging being the main activity in flocks, birds in flocks frequently vocaliseto an audience of both heterospecific and conspecific participants (Goodale & Kotagama 2008, Pagani-Núñez *et al*. 2018, Coppinger *et al*. 2020). For example, alarm vocalizations constitute an anti-predatory strategy, which enables the recruitment of other species for mobbing and reduces predation risk (Goodale & Kotagama 2008, Fallow *et al*. 2013, Magrath *et al*. 2020, Zhou *et al*. 2021). In parallel, participant species may eavesdrop on other species’ vocalizations, use them to assess predation risk, and signal to heterospecifics (Goodale *et al*. 2017). In the crowded milieu of a flock, this diversity of sounds may result in competition for acoustic space due to interference between signals (Bee & Micheyl 2008). Additionally, morphological traits influence the properties of emitted sounds (Gillooly and Ophir 2010), and it is therefore important to examine whether the two sets of traits exhibit similar community-level patterns (Krishnan and Tamma 2016). However, community-scale approaches that quantify the trait space and vocal activity patterns of participant species in flocks are lacking, and may inform us about their role in flock assembly and communication.

Here, we examine the organization of acoustic signal trait space of participant species in flocks in relation to their morphological trait space, employing a community phylogenetic approach (Cavender-Bares et al, 2004). These patterns are examined across flock sizes, because the benefits of participating in flocks may vary with flock size (Bangal *et al*. 2022). Many mixed-species flocks across the world are known to consist of similar-sized participant species (Sridhar *et al*. 2012), which benefit from an enhanced dilution effect when encountering a predator. Body size exerts effects on acoustic signals — for example, larger species exhibit lower frequencies (Wallschläger 1980, Goller & Riede 2013, Liu *et al*. 2017) — and we hypothesize that this effect may drive similarity in acoustic trait space as a by-product (Krishnan & Tamma 2016).

Studying mixed-species flocks of peninsular Indian deciduous forests, we addressed the following questions: a) Do these flocks consist of similar-sized species? b) Does morphological similarity drive similarity in acoustic trait space? Alternatively, if acoustic trait space is structured by the need to communicate effectively, we predicted a pattern of dissimilarity in acoustic signal trait space in flocks (Cavender-Bares *et al*. 2004, Chhaya *et al*. 2021). This would enable participant species to maximize the benefits of being similar in body mass and simultaneously minimize the cost of masking interference owing to acoustic similarity (Bee & Micheyl 2008). Dissimilarity in acoustic signals could alternatively arise from phylogenetic non-independence, i.e., the flocks being composed of distant relatives which are therefore also different in their acoustic signals. Thus, we also performed phylogenetically informed analyses to disentangle these possibilities.

## Materials and Methods

### Data collection

We conducted the study in Wayanad Wildlife Sanctuary (WWS), a moist-deciduous forest located in the north-eastern region of Kerala, India, following transect sampling to collect data on mixed-species flocks. We collected data across a total of 39 transects (2 km each with a gap of at least 250 m between them), walking each transect twice between December 2021 and April 2022. Two sets of data were collected for each encountered flock, which directly informed the analyses:

#### Acoustic recordings of flocks

Vocalizations of flocking birds were recorded using a combination of a Sennheiser (Wedemark, Germany) ME-66 directional microphone with a K6 power module and a Zoom (Tokyo, Japan) H6 recorder. Vocalizations were all recorded at a sampling rate of 48 kHz with 16-bit resolution and were saved in .*wav* format. We used the following criteria to identify a flock: 1) Presence of an active foraging group of birds, 2) the group should have at least two or more species, and 3) the group should be moving in the same direction for a minimum of two minutes. After this two-minute window, we recorded the vocalizations in the flock for aanywhere between 5 and 15 minutes, depending on how long we were able to follow the flock. In total, we obtained acoustic data from 112 flocks.

Because the distance between the microphone and vocalizing birds can significantly impact the received amplitude, and thus the quality of the recordings, we obtained all recordings within a maximum of 30 metres distance from the flock. Flocks are not stationary and to maintain this distance, we followed the flock as it was being recorded.

#### Visual census of flock participation

Throughout the observation time of each encountered flock, we also made note of all species present in the flock. An individual of any species was considered part of the flock if it was detected visually within a 10 meter radius of its nearest neighbour in the group. For consistency in data entry, we used the following criteria: 1) Only individuals that were detected visually within the above radius were considered flock participants, regardless of whether their calls were recorded or not. 2) Only species that maintained their participation in the flock throughout the recording duration were considered participant species of the flock. Although conservative, this ensured a consistent “participant list” that could then be matched to the acoustic data, as it was impossible to identify flock participants from acoustic recordings alone. The data collected on the species composition of flocks were tabulated during the field data collection phase. This is henceforth referred to as *the visual census data* on the species composition of flocks.

### Analysis

The extraction of information from the sound recordings (hereafter referred to as the *acoustic census data*) of vocal participants in the flocks was carried out separately using the Raven Pro 1.6 software (Cornell Laboratory of Ornithology, Ithaca, NY, USA). We followed a dispersed sub-sampling method (Supplementary Figure 1) to tabulate acoustic census data of species from the recordings. For this, we sampled every alternate minute of each recording using species composition data from the visual census as a reference, and made note of all vocal species in each time sample.

Acoustic census data was converted to a Presence-Absence (PA) Matrix for analysis, where a value of 1 meant that the species’ vocalization was detected (regardless of the call type) and a value of 0 meant the absence of any vocalization of that species. A very small number (< 1%) of vocalizations could not be identified and were not included in the analysis.

#### Categorization of visual and acoustic census data

Using the visual census matrix, we identified three flock species richness categories. Flocks with species richness between 2 and 5 were categorized as small, flocks with species richness between 6 and 10 were categorized as medium, and flocks with more than 10 species were categorized in the large flock richness category, following criteria laid out in published studies (Bangal *et al*. 2021).

For the acoustic census matrix, there were no flocks with >10 vocalizing species in any one-minute time frame. Therefore, we categorized flocks into two vocal activity categories. Time frames with 2-5 vocal species were categorized as low vocal activity, and time frames with 6-10 vocal species were categorized in the high vocal activity category. The flock richness and the vocal activity categories are used in the subsequent analysis detailed below.

#### Vocal activity of flock participants

We calculated a vocal activity index for each species as the proportion of time frames in which a participant species was acoustically detected in the recordings, (i.e., the number of time frames with values 1 divided by the total number of time frames in which the species was present in flocks, as determined by visual census).

#### Acoustic signal parameters of participant species

Apart from the vocal activity of participant species, we investigated the distribution of species within the acoustic signal space of flocks. For this purpose, we calculated signal parameters for each participant species using Raven Pro. Spectrograms were generated using a Hann window with a sample size of 512 samples and a 50% overlap between windows. For each species, we labelled at least 20 notes using the cursor and selection box features in Raven Pro. While digitizing the notes, we took care to encompass the diversity of the vocal repertoire for each species that was recorded. To ensure this, we digitized all notes in a species’ call/song. In the case of relatively vocal species, different notes in the repertoire were digitized in similar/comparable proportions to avoid over-representation of a particular note.

We calculated the following eight parameters for all the digitized notes: Peak Frequency (Hz), the first, last, maximum and minimum Frequencies of the Peak Frequency Contour (Hz) (using the peak frequency contour feature in Raven Pro), Average Entropy (bits), Centre Frequency (Hz), and Bandwidth containing 90% of energy (Hz). These parameters were chosen as they are relatively robust to variation in amplitude and subjectivity in note digitization across the recordings (Chitnis *et al*. 2020, Lahiri *et al*. 2021). For the acoustic variables, we performed a Principal Components Analysis on the correlation matrix of eight acoustic variables in R. Because the first principal component explained the most (∼75 %) variation (Supplementary Table 2), we used the mean PC1 score for each species to test for acoustic phenotypic patterns in subsequent analyses.

#### Correlation between body mass and vocal activity index of participant species

Finally, we examined the relationship between body mass (sourced from AVONET, a bird morphometric dataset (Tobias 2022)) and vocal activity of participant species (that participated in more than 5% of the flocks). We calculated Spearman’s correlation coefficient, a rank order correlation used for non-normal data (the body mass of the species was not normally distributed) (Supplementary Figure 2). This was used to quantify the direction and the strength of association between body mass and vocal activity index of participant species. Comparative analyses of traits run the risk of increased Type I errors owing to phylogenetic non-independence between species (Felsenstein 1985, Revell *et al*. 2008). Therefore, we transformed the two variables into phylogenetically independent contrasts (PIC) using functions in the *ape* (Paradis *et al*. 2019) package in R version 4.2.0 (R Core Team, 2022). The phylogenetic trees required for this analysis were sourced from the Bird Tree of Life project (http://birdtree.org/) (Jetz *et al*. 2012), and were pruned to include only the species in our dataset. A consensus tree was generated from the 1000 possible phylogenetic hypotheses, using the consensus.edges() function and least squares method in the *phytools* package (Revell and Revell, 2014) in R. The transformed variables were then used to estimate and test the correlation between body mass and vocal activity index of flock participants, to examine whether any underlying correlation of these variables was influenced by the phylogenetic relatedness of species.

#### Phenotypic similarity analysis

To understand phenotypic assembly in flocks of different richness and vocal activity categories, we examined morphological (body mass) and acoustic (PC1 of acoustic parameters) similarity/overdispersion in flocks. We calculated the standard deviation of the trait values for each flock as a measure of flock level variation in the phenotypic trait of interest (Bangal *et al*. 2021). These observed values of variation were averaged across each richness (small, medium and large) and vocal activity (low and high) category. Here, a small value of standard deviation indicated phenotypic similarity and a large value indicated overdispersion. To understand if the observed phenotypic variation at different flock richness and vocal activity categories differed from chance expectations, we generated null flocks using both visual and acoustic census matrices. First, we randomized the species-by-flock matrices (both acoustic and visual matrices) 1,000 times, maintaining the flock sizes (column total) as constant and assigning probabilities of occurrence to each species based on the observed values of occurrence (Gotelli 2000). The null flocks generated from the visual and acoustic census matrices were also divided into three flock richness categories and two vocal activity categories, respectively, as detailed above.

Null models were generated using the sim5 function in the *EcoSimR* package (Gotelli *et al*. 2015). This algorithm inputs a species-by-flock matrix and assigns weights to each species based on the probability of occurrence in the entire dataset. Therefore, when generating null flocks, the function draws each species based on the probability of a particular species participating in a flock. Therefore, the columns in an empty matrix were filled proportional to the row sums (species occurrence), simultaneously holding the column totals as fixed values (i.e., the flock richness is kept constant). During each simulation, the average standard deviation of trait values (body mass and acoustic variables) was calculated within richness categories for each category. This null model approach thus controlled for the effect of flock size on the average values of the acoustic and morphological traits. To compare null models to the observed data, we calculated a Standardized Effect Size (SES).

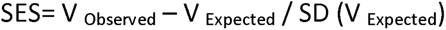

Where V _Observed_ is the observed variation of the mean trait values, V _Expected_ is the mean obtained from 1,000 randomized simulations and SD (V _Expected_) is the standard deviation around the mean trait values from the simulated data. A positive SES indicates a larger effect than expected by chance, and a negative value indicates the converse, i.e. less variation than expected by chance. Statistically, values less than -1.96 and greater than 1.96 are considered significant as the values of the standardized effect size form a standard normal distribution and 2.5% of the observations lie above and below 1.96 and -1.96 respectively (Gotelli & McCabe 2002).

#### Phylogenetically corrected analysis for traits

To measure phylogenetic signal, we calculated Blomberg’s *K* (Blomberg *et al*. 2003) for body mass and PC1 of the acoustic parameter space, using the tree obtained earlier (Revell and Revell, 2014). The significance (*p-value*) of *K* was determined by comparing the observed *K* value to the null distribution of *K* values calculated from 1000 randomized trees, generated by shuffling the tips of the tree used to calculate the observed *K*.

We additionally examined whether trait space patterns were influenced by phylogenetic signal using Faith’s phylogenetic diversity (PD) for each observed flock, calculated using the *picante* package (Kembel et al. 2010) in R. Faith’s PD is calculated by summing the branch lengths of the tree, pruned for each observed flock. Because PD is correlated with species richness (Cadotte et al. 2011), we generated null flocks to assess whether the observed PD was significantly different from the values expected by chance. For this, we generated null distributions by randomising the visual census matrix 999 times using the ‘independent swap’ method (Gotelli & McCabe, 2002, Gotelli & Entsminger, 2003). This method preserved values of species richness and the number of flocks in which each species was found. We further calculated the SES values for each flock, a measure of the difference between the observed PD and the mean of the null PD generated for each flock.

## Results

### Flocks with small-to-medium species richness consist of similar-sized species

We found that species participating in small and medium richness flocks (i.e., 2-10 species per flock, constituting ∼87% of total flocks observed) were significantly more similar (lower standard deviation) in body mass than expected by chance (Figure 1a: SES= -3.90, p < 0.05 for small-flock richness category, SES= -3.72, p < 0.05 for medium-flock richness category). Among these two categories, small-richness flocks exhibited greater similarity in body mass than medium-richness flocks (Figure 1a). In the large-richness flocks (13% of total flocks), however, the observed variation in body mass among participant species lay within the expectation under chance (SES= 1.41, p > 0.05) (Figure 1a). Body mass distributions across the three flock richness categories suggested that species with smaller body masses dominate across all flocks (Figures 2a and 2b), but particularly small and medium-richness flocks.

**Figure 1.**
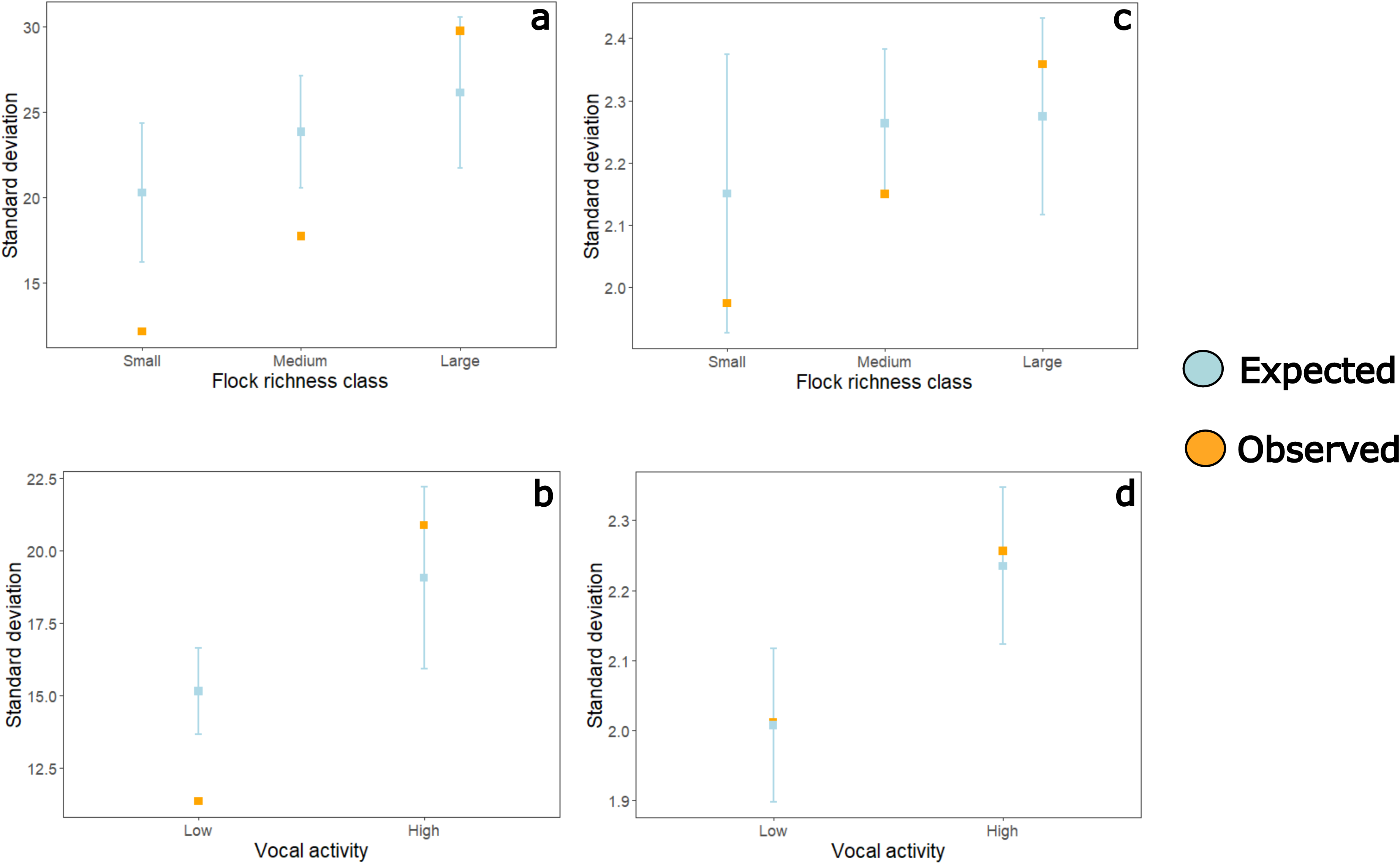
Observed (orange) and expected (blue) variation of body mass across **a.** small-, medium-, and large-flock richness categories as determined by visual census and **b.** low and high vocal activity categories, as determined from acoustic census. Observed and expected variation in acoustic PC1 across **c.** small-, medium-, and large-flock richness categories, as determined by visual census and **d.** low and high vocal activity categories, as determined by acoustic census. The orange points are the observed standard deviation of participant species’ body mass averaged over flocks in each richness category. The blue points are the average of the standard deviation of body mass from 1000 simulations (null flocks) and the blue lines represent 2*standard deviation of the variation in body mass generated from the 1000 simulations for each flock richness category.

**Figure 2.**
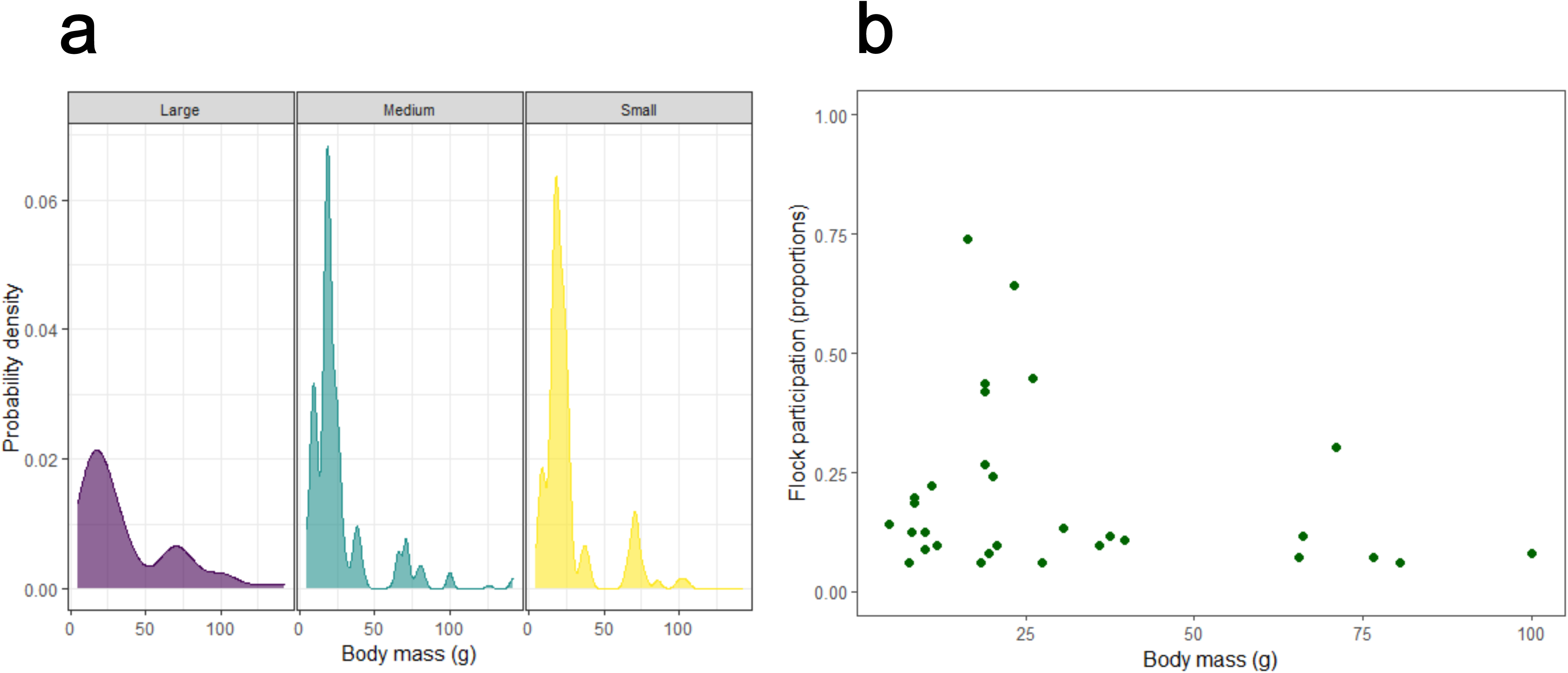
a. Smoothed histogram of body mass of participant species in three flock richness categories**. b.** The relationship between frequency of flock participation (proportion of flocks in which each species was observed) and body mass of species. Species observed in more than five percent of flocks are depicted here, representing the community of flock-participating species.

### Body mass similarity among species that temporally vocalise together

Next, we compared patterns in body mass for species that were detected in our acoustic census, i.e., species that are vocal participants of mixed-species flocks (grouped into categories based on how many species were vocal). Similar to the visual census, we found that species that were acoustically detected in flocks with low vocal activity (2-5 species vocalising together within a sampling time frame) were more similar in body mass than expected by chance (Figure 1b: SES= -4.86, p < 0.05). Approximately 86% of all the analysed time frames represented flocks with low vocal activity. However, this similarity pattern did not hold for species present in the high vocal activity category (SES= 1.16, p > 0.05) (Figure 1b).

### Smaller participants tend to vocalise more while foraging in flocks

Next, we examined the relationship between body mass and vocal activity index of participant species in flocks, as a prelude to examining the acoustic trait space. The vocal activity index (as determined from the acoustic census data) was negatively correlated with body mass of the flock participants (Figure 3a, Spearman’s correlation coefficient = -0.47, p = 0.009). Larger (Figure 4a) and less vocal species (Figure 4b) were also phylogenetically closely related. Consistent with this, the correlation between the two variables was not statistically significant in a phylogenetically corrected analysis (Figure 3b, Spearman correlation coefficient= -0.25, p= 0.2).

**Figure 3.**
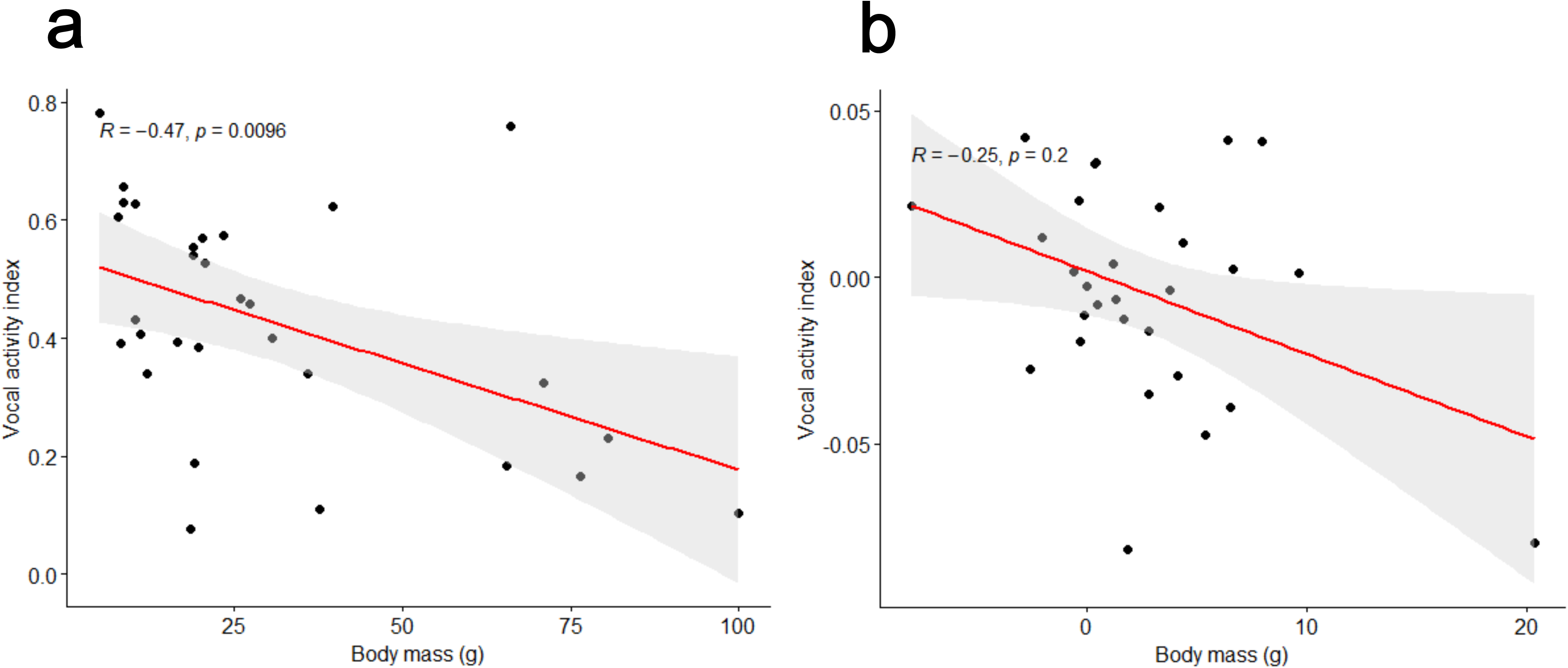
a. The relationship between vocal activity index and body mass of participant species in flocks. Data points represent species that were observed in >5 % of sampling time frames. **b.** The relationship between vocal activity index and body mass is not statistically significant after correcting for phylogenetic non-independence.

**Figure 4.**
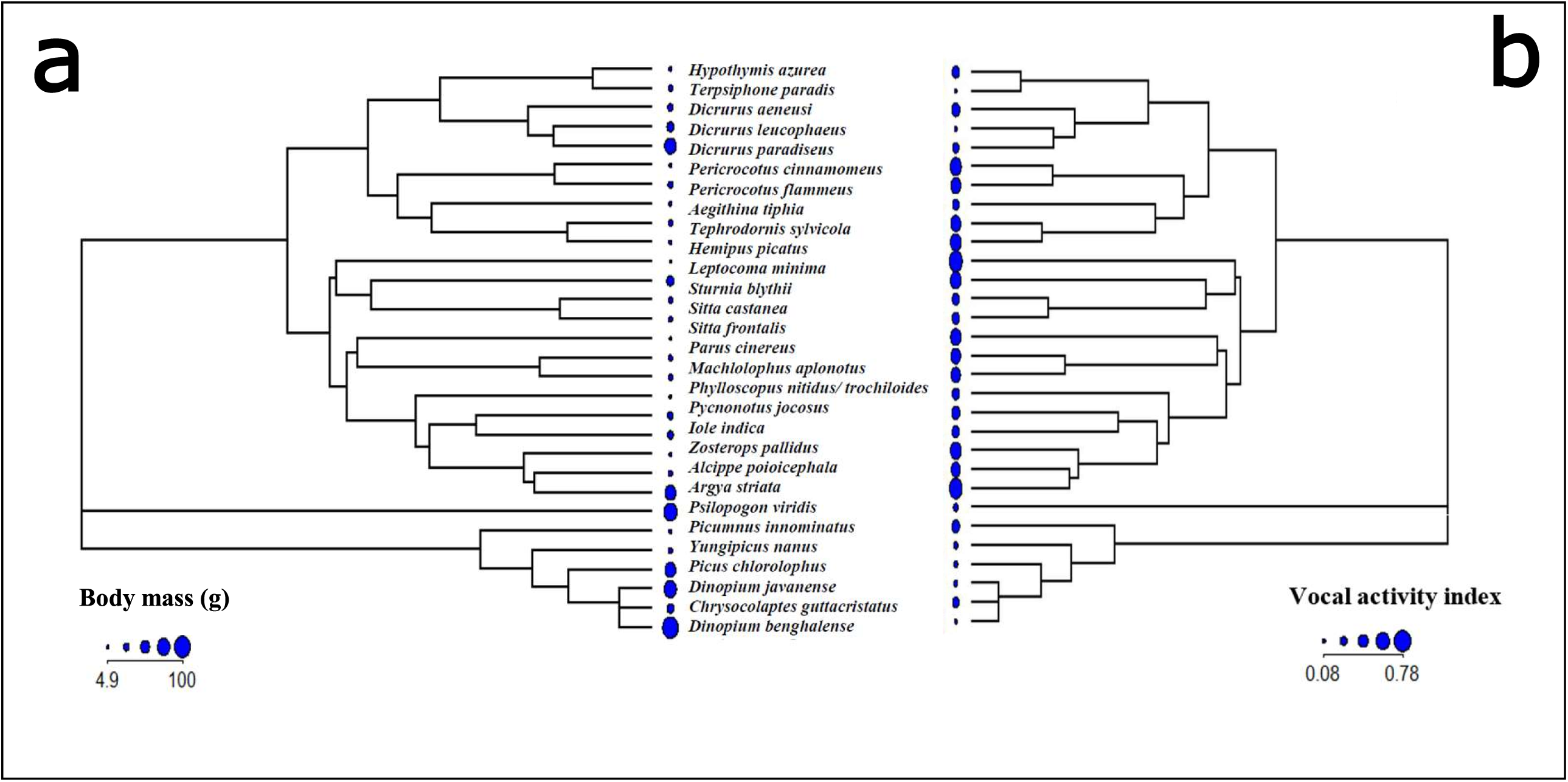
a. Consensus of 1000 phylogenetic trees representing the relationships of species participating in flocks. The blue circles represent body mass of participant species, and larger size indicates higher mass. Note that larger species are clustered together on the tree. **b**. Consensus of 1000 phylogenetic trees representing the relationships of species participating in flocks. The blue circles represent vocal activity index of participant species, and larger size indicates higher vocal activity. Note that vocally active species are also clustered on the tree.

### Mixed species flock participants are not similar in acoustic parameter space

Our data up to this point suggested that flocks are highly similar in body mass, particularly flocks with lower species richness, and that they tend to consist of smaller species that are highly vocal (within flocks), this emerging as a consequence of being relatively closely related. In a Principal Components Analysis (PCA) that summarised eight acoustic variables (see Methods), the first two Principal Components (PC’s) accounted for 95% of the total variation (Supplementary Table 2). All six parameters loaded negatively on PC1, whereas average entropy and bandwidth loaded negatively on PC2. The first principal component accounted for ∼75% of the variation in data. Using the average PC1 values for each species, we found that acoustic parameters were not significantly similar for participant species across all flock richness categories (Figure 1c: SES= -1.49, p > 0.05 (small-flock richness category), SES= -1.93, p > 0.05 (medium-flock richness category), SES= 1.08, p > 0.05 (large-flock richness category)). Unlike the body mass distribution of species, the distributions of acoustic PC1 across the three flock richness categories were not skewed (Supplementary Figure 3).

Similar patterns held for species vocalizing together across all time frames, when we considered participation using categories derived from the acoustic census matrix. Regardless of which census method was used, participating species were not statistically similar in acoustic signal space. SES values were within the range expected under chance (Figure 1d: SES= 0.16, p > 0.05 for the low-vocal activity category, SES= 0.36, p > 0.05 for the high-vocal activity category).

### Phylogenetic signal of body mass and acoustic PC1 of participant species

Blomberg’s *K* for both the body mass and the acoustic traits of participant species indicated significant phylogenetic signal (body mass: *p* < 0.05, acoustic PC1: *p* < 0.05). K values for both the body mass and acoustic PC1 were 0.4, suggesting that closely related species were more divergent in trait values than expected under Brownian motion.

Because both morphology and acoustic signal traits exhibited significant phylogenetic signal, we calculated Faith’s PD for each flock and compared the values with null flocks. The distribution of SES values for all flocks (Figure 5) indicates no departure from chance, and thus a lack of phylogenetic clustering. Therefore, the observed morphological similarity, and the patterns in acoustic space, were not influenced by phylogenetic clustering within flocks.

**Figure 5.**
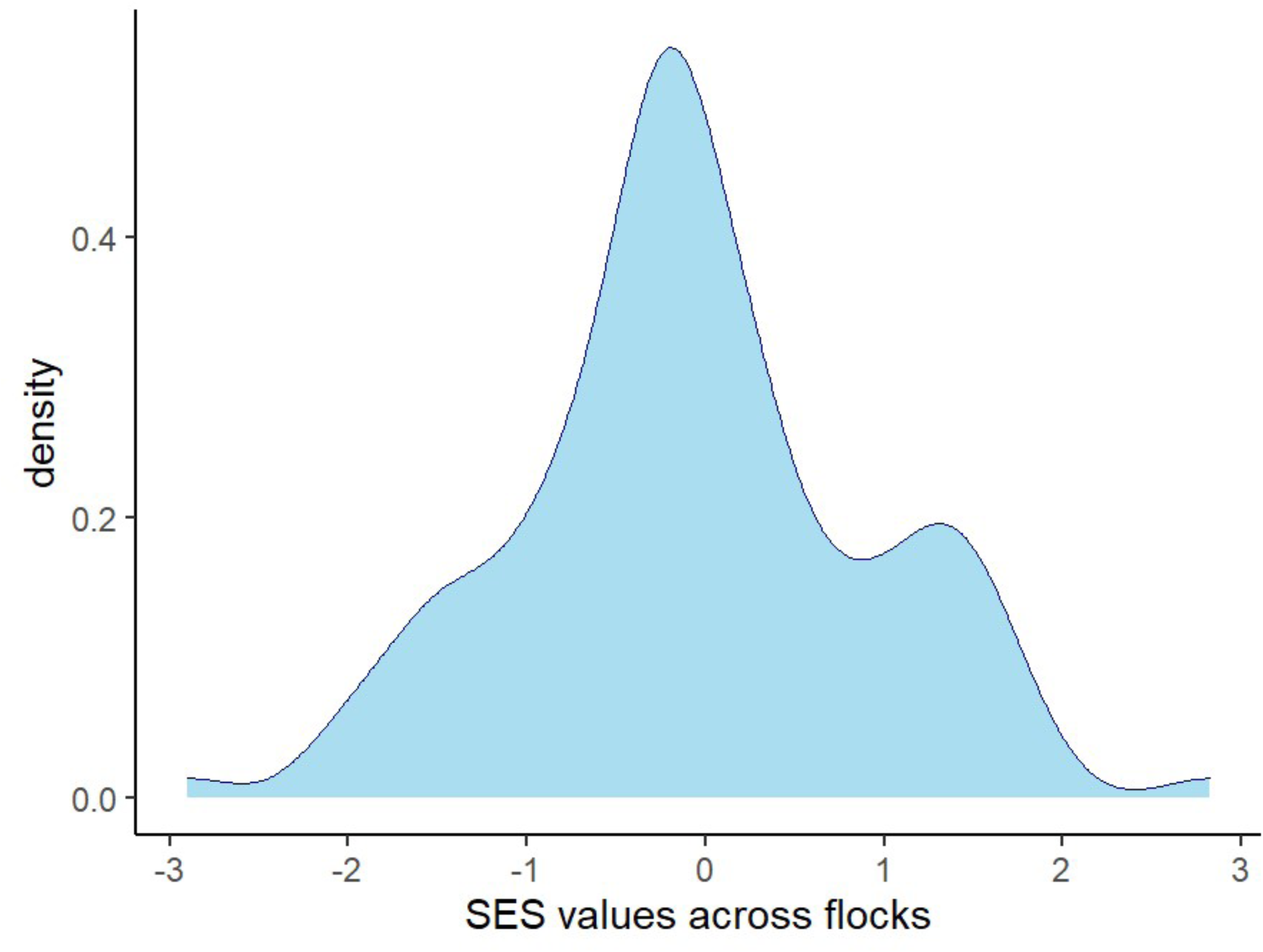
Smoothed histogram of SES values generated from Faith’s PD values calculated for observed and null flocks.

## Discussion

Our study found that participants in mixed-species flocks are more similar in body mass than expected by chance. This body mass similarity pattern was observed among species both in small and medium flock richness categories (∼87% of observed flocks), but not in large flocks. The trade-offs between the benefits and costs associated with joining small versus large flocks are discussed in the literature, and our findings corroborate these (Bangal *et al*. 2021). For instance, large flocks open different niche spaces and thereby accommodate participants that vary in morphology and different foraging behaviours (Harrison & Whitehouse 2011) whereas in small flocks, benefits are related to being similar in body size, and hence increased dilution of risk, among other benefits.

In addition, we examined the body mass similarity patterns across vocal activity categories, which represent a measure of *vocal* participation in flocks as opposed to the visual census data discussed above. The Low vocal activity category constituted ∼86% of all analyzed time frames in the acoustic census, and these vocalizing species were also more similar in body mass than expected by chance i.e., smaller-sized birds which dominated flocks were also the most vocal. Larger species such as woodpeckers and the Greater Racket-tailed drongo (*Dicrurus paradiseus*) vocalised less frequently when participating in flocks. The sole exception to this was the Jungle Babbler (*Argya striata*), which was the only large-bodied, gregarious species that was also very vocal in flocks. Broadly, smaller-sized and highly vocal species were also relatively closely related to each other on a phylogeny, and so the similarity in vocal activity appears to be a consequence of their phylogenetic relatedness. Would this similarity in body mass, and proximity on a phylogeny, therefore also translate to overlap in acoustic signal space? Contrary to this expectation, we found that flock participants (which were similar in body mass) were not similar in acoustic signal characteristics. A comparison between observed and expected acoustic signal traits across all flock richness and vocal activity categories (determined using both acoustic and visual census methods) revealed no departures from the expectation under chance. This pattern, of similarity in morphological but not acoustic trait space, did not appear to be influenced by phylogenetic relatedness. In support of this, we did not find evidence of phylogenetic clustering in flock participation even though the traits themselves exhibited significant phylogenetic signal.

Morphology and acoustic signal parameters are known to be correlated (Gillooly & Ophir 2010), however, results from this study provide initial evidence to suggest that the two trait spaces are decoupled at the level of multispecies associations. This decoupling suggests that participant species in flocks may potentially exhibit similarity in body mass without overlap in acoustic signal parameters, and are thereby expected to derive distinct sets of benefits from each. Similar-sized individuals may experience similar predation pressures (Gliwicz 2008), and thereby benefit from the dilution effect when participating in flocks. On the other hand, possessing distinct acoustic signals supports interspecific signalling (Magrath *et al*. 2015, 2020), sharing social information (Goodale *et al*. 2010), and activity matching (Sridhar & Guttal 2018) between species. Most acoustically signalling animals communicate in crowded conditions, where signals propagate over a host of other biotic and abiotic sounds. Because an acoustic community comprises multiple signallers both within and across taxa, their vocalizations may overlap in acoustic space, further reducing signal efficacy and leading to masking interference (Schwartz & Wells 1983, Bee & Micheyl 2008, Balakrishnan *et al*. 2014). Processes such as these may drive community-level patterns ranging from divergence (overdispersion) to convergence (similarity) of acoustic parameters in the signal space (Luther 2009, Krishnan 2019). Combining behavioural studies with an analysis of acoustic trait space thus helps examine which processes drive community-level patterns (example, Sugai *et al*. 2021). Competition for signal space is predicted to result in the divergence of acoustic parameters, whereas acoustic convergence is predicted in extended interspecific communication networks (Tobias *et al*. 2014).

In light of the conceptual framework discussed above, community-level trait patterns offer insights into species assembly within flocks. Mixed-species groups present an opportunity for species to vary along different trait axes, unlike single-species groups where traits are expected to be correlated. For example, Sridhar & Guttal (2018) showed that participant species exhibit decoupling between body mass and foraging behaviour i.e., species with same body masses might have strong associations in flocks, but exhibit different foraging behaviours. This enables birds to reduce competition along a certain trait axis (foraging behaviour in this example), and simultaneously benefit from similarity in morphological traits (which dilutes the risk of being predated).

Our study observes a similar decoupling at the community level between morphological and acoustic traits in these transient interspecific associations. The opposite pattern is observed in sympatric communities of closely-related Asian Barbets, where the morphological and acoustic traits of congeneric sympatric barbets are apparently coupled (Krishnan & Tamma 2016). In flocks, however, which are a dynamic mix of closely and distantly related species, decoupling of the acoustic and body mass trait axes potentially maximizes the benefits of being similar in body mass, and yet minimizes the costs of being acoustically similar. Here, it is important to emphasize that similar results hold when investigating trait patterns using different census and observation methods.

Simultaneously, it is also important to note that even though species are not similar in acoustic space, they are also not overdispersed (i.e., they are not *more* divergent than expected by chance). The divergence of acoustic trait space is predicted under situations where the priority is for species to signal to conspecifics alone. In the context of sexual signals, for instance, heterospecific signals are a potential source of interference or noise (Goodale *et al*. 2019). However, similarity between signals impedes the segregation of sound sources (MacDougall-Shackleton *et al*. 1998, Nityananda & Bee 2011), and creates confusion among predators (Goodale *et al*. 2019), especially when multiple prey individuals are signalling simultaneously. In other words, it pays to be distinct enough to be recognized by potential mates (conspecifics) and flocking companions (heterospecifics), and at the same time have some overlap in acoustic signal space to confuse predators. The patterns we found in the vocal activity within flocks also lend support to this argument. Therefore, we hypothesize that the observed acoustic trait space (neither similar nor overdispersed) among species in flocks concurrently provides the potential benefits of reducing masking interference within the flock, enhancing interspecific communication (non-alarm), and leaving room for some convergence in vocalizations, putatively to confuse predators. However, the acoustic census matrix was tabulated based on the vocal detection of a species, irrespective of the number of individuals or behavioural context. Therefore, pending further research on the contexts of vocalizations (such as alarm calling, contact calling, etc.), our results point the way to future playback experiments and detailed examination of the functional basis of acoustic communication in flocks.

## Acknowledgments and funding

We thank the NCBS-TIFR, JNCASR and the Nature Conservation Foundation for institutional, funding, and administrative support. AA would like to thank Mathan Pilakkavu, Manoj Kumar A V, N Badusha, and Arul Badsha for their support during fieldwork. We are also thankful to the Kerala Forest Department for issuing permits to conduct this study within Wayanad Wildlife Sanctuary (Permit number: KF DH Q - 42331202 1 .CWW/WLI 0). A.K. is funded by a core research grant (Science and Engineering Board (SERB), Government of India (CRG/2022/000187)). This study was approved by the NCBS-Institutional Animal Ethics Committee (proposal Number: NCBS_IAE_2021/ 07(N)). We declare we have no competing interests.

## Figure legends

**Supplementary Figure 1.** A representation of dispersed sub-sampling method for a 7-minute audio recording.

**Supplementary Figure 2.** Smoothed histogram of body mass of participant species

**Supplementary Figure 3.** Smoothed histogram of acoustic PC1 of participant species in three flock richness categories

**Supplementary Figure 4.** Dendrogram representing similarity in species composition between flocks observed at different locations in Wayanad Wildlife Sanctuary, Kerala. Similarity between flocks was calculated using the Euclidean distance metric where flocks that are similar in species composition group together. The dendrogram is built using hierarchical clustering and the complete linkage method. Each terminal node represents an individual flock and is colour-coded based on the location (Field base camps) where it was observed i.e., flocks color-coded by the same colour were observed at the same location.

## Tables

**Supplementary Table 1.** Species and their corresponding flock participation (in percentage)

| *Species* | *Percentage of occurrence in flocks (%)* |
| --- | --- |
| *Yungipicus nanus* | 26.8 |
| *Tephrodornis sylvicola* | 24.1 |
| *Hypothymis azurea* | 22.3 |
| *Zosterops palpebrosus* | 19.6 |
| *Pericrocotus cinnamomeus* | 18.8 |
| *Leptocoma minima* | 14.3 |
| *Iole indica* | 13.4 |
| *Hemipus picatus* | 12.5 |
| *Phylloscopus nitidus/ trochiloides* | 12.5 |
| *Dicrurus leucophaeus* | 11.6 |
| *Argya striata* | 11.6 |
| *Sturnia blythii* | 10.7 |
| *Alcippe poioicephala* | 9.8 |
| *Aegithina tiphia* | 9.8 |
| *Chloropsis aurifrons* | 9.8 |
| *Picumnus innominatus* | 8.9 |
| *Sitta castanea* | 8 |
| *Dinopium benghalense* | 8 |
| *Dinopium javanense* | 7.1 |
| *Picus chlorolophus* | 7.1 |
| *Culicicapa ceylonensis* | 7.1 |
| *Terpsiphone paradisi* | 7.1 |
| *Pycnonotus jocosus* | 6.3 |
| *Psilopogon viridis* | 6.3 |
| *Chrysocolaptes guttacristatus* | 4.5 |
| *Dendrocitta vagabunda* | 4.5 |
| *Dicrurus caerulescens* | 4.5 |
| *Hemicircus canente* | 3.6 |
| *Acridotheres fuscus* | 3.6 |
| *Loriculus vernalis* | 3.6 |
| *Irena puella* | 2.7 |
| *Gracula indica* | 2.7 |
| *Coracina macei* | 2.7 |
| *Harpactes fasciatus* | 2.7 |
| *Cinnyris asiaticus* | 2.7 |
| *Cyornis tickelliae* | 2.7 |
| *Eumyias thalassinus* | 2.7 |
| *Lalage melanoptera* | 1.8 |
| *Brachypodius priocephalus* | 1.8 |
| *Oriolus kundoo* | 1.8 |
| *Phylloscopus magnirostris* | 1.8 |
| *Pycnonotus cafer* | 1.8 |
| *Orthotomus sutorius* | 1.8 |
| *Leiopicus mahrattensis* | 1.8 |
| *Muscicapa dauurica* | 0.9 |
| *Turdus simillimus* | 0.9 |
| *Dumetia atriceps* | 0.9 |
| *Oriolus chinensis* | 0.9 |
| *Merops leschenaulti* | 0.9 |
| *Argya subrufa* | 0.9 |
| *Psittacula columboides* | 0.9 |
| *Dicaeum concolor* | 0.9 |
| *Psittacula cyanocephala* | 0.9 |
| *Pellorneum ruficeps* | 0.9 |
| *Rubigula gularis* | 0.9 |
| *Micropternus brachyurus* | 0.9 |
| *Arachnothera longirostra* | 0.9 |

**Supplementary Table 2.**
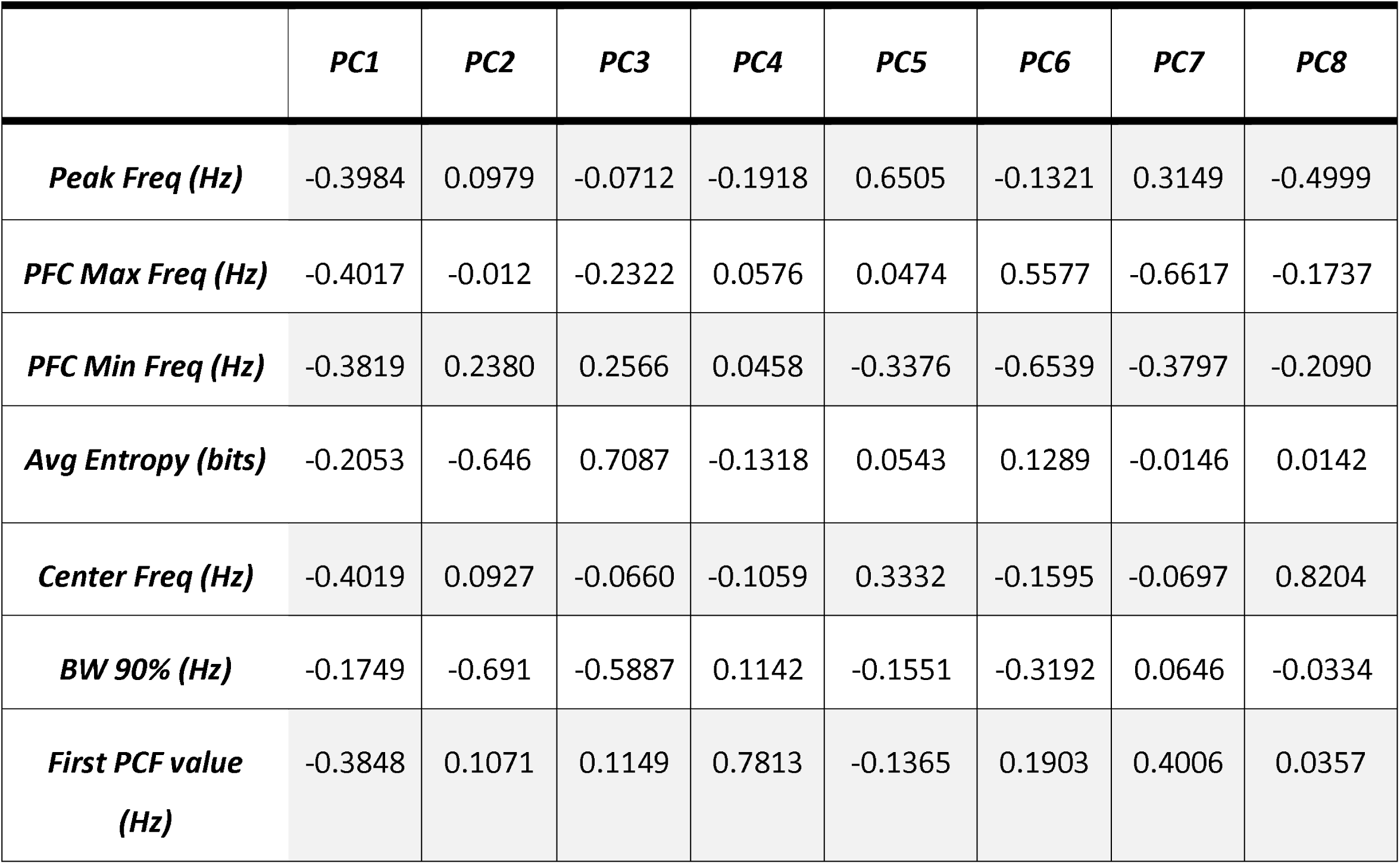
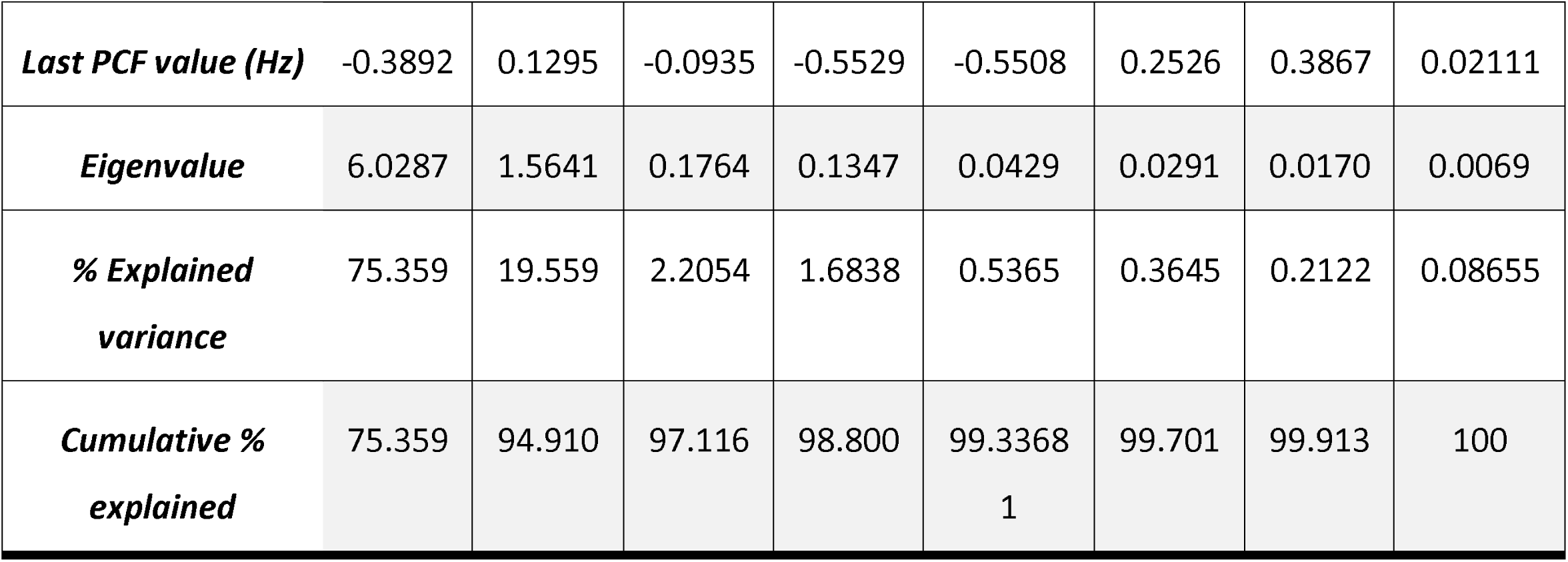
Results of principal component analysis for eight acoustic variables. Rows contain factor loadings for each trait, eigenvalues and the proportion of variance explained by each principal component.

