## Supplementary Material for "Contrasting morphological and acoustic trait spaces suggest distinct benefits to participants in mixed-species bird flocks"

**Supplementary methodology**

*Examining flock composition similarity across field base camps*

We calculated Euclidean distances between flocks based on their composition as determined from 78 walks of 39 transects and constructed dendrograms using hierarchical clustering and complete linkage (base R functions dist() and hclust()) to visualize location-specific differences in flock composition (R core Team 2022).

**Supplementary result**

*Flock composition is not location-specific*

Prior to studying general flocking and phenotypic patterns of birds across the study area, it was imperative to determine whether habitat characteristics or sampling location influenced the flock composition. Cluster analysis on flocks based on their composition revealed that flocks from the same location did not cluster together (Supplementary Figure 4). In the dendrogram, each node represents a flock, and is color-coded based on the area around which it was observed. Because the colours representing different sampling areas were dispersed across the dendrogram without clustering, we thus inferred that flock composition did not exhibit location-specific patterns, with similar compositions across the study area.
