## Supplementary figures and images for "Contrasting morphological and acoustic trait spaces suggest distinct benefits to participants in mixed-species bird flocks"

### Supplementary Figure 1

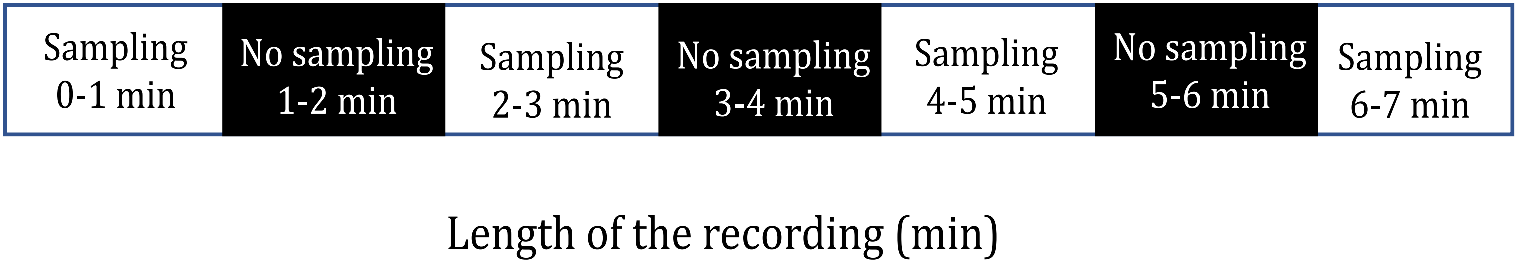

### Supplementary Figure 2

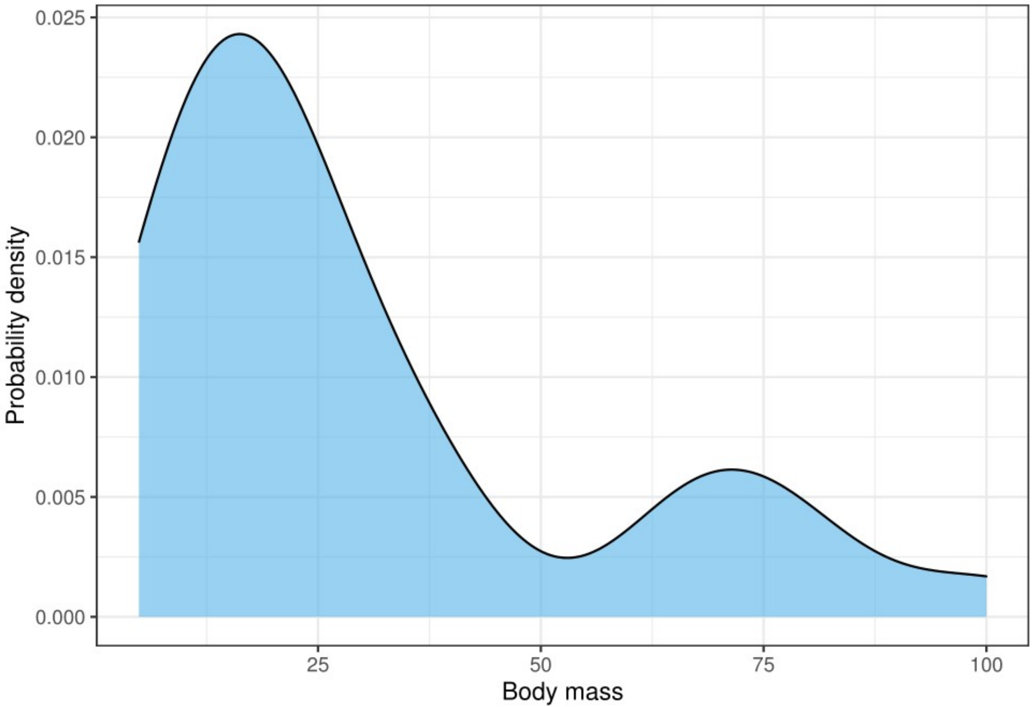

### Supplementary Figure 3

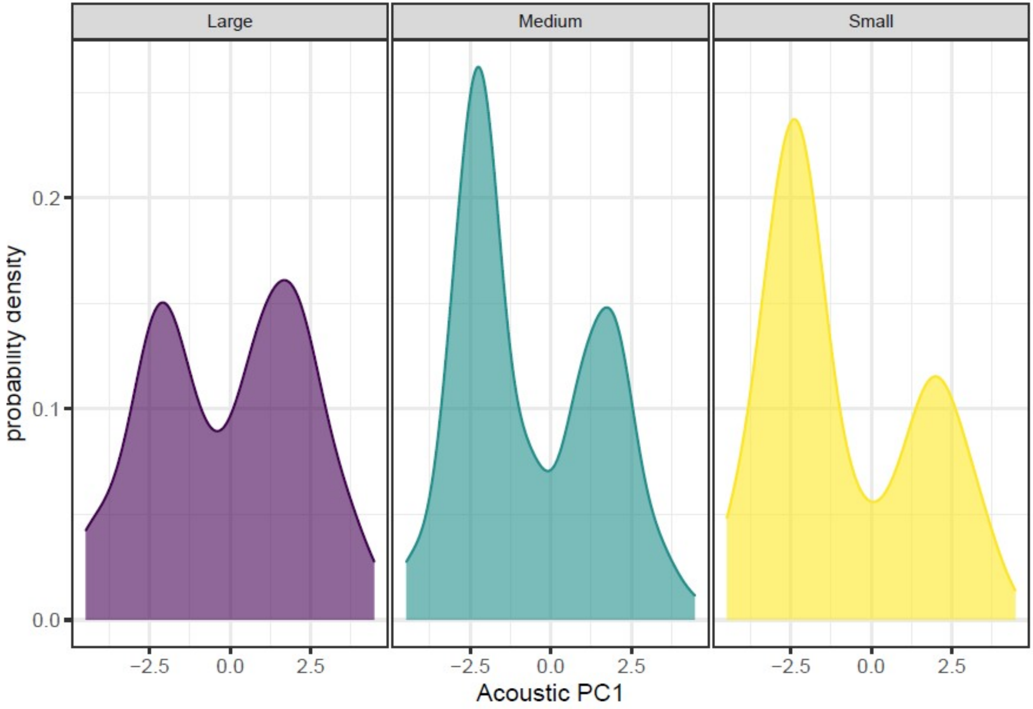

### Supplementary Figure 4

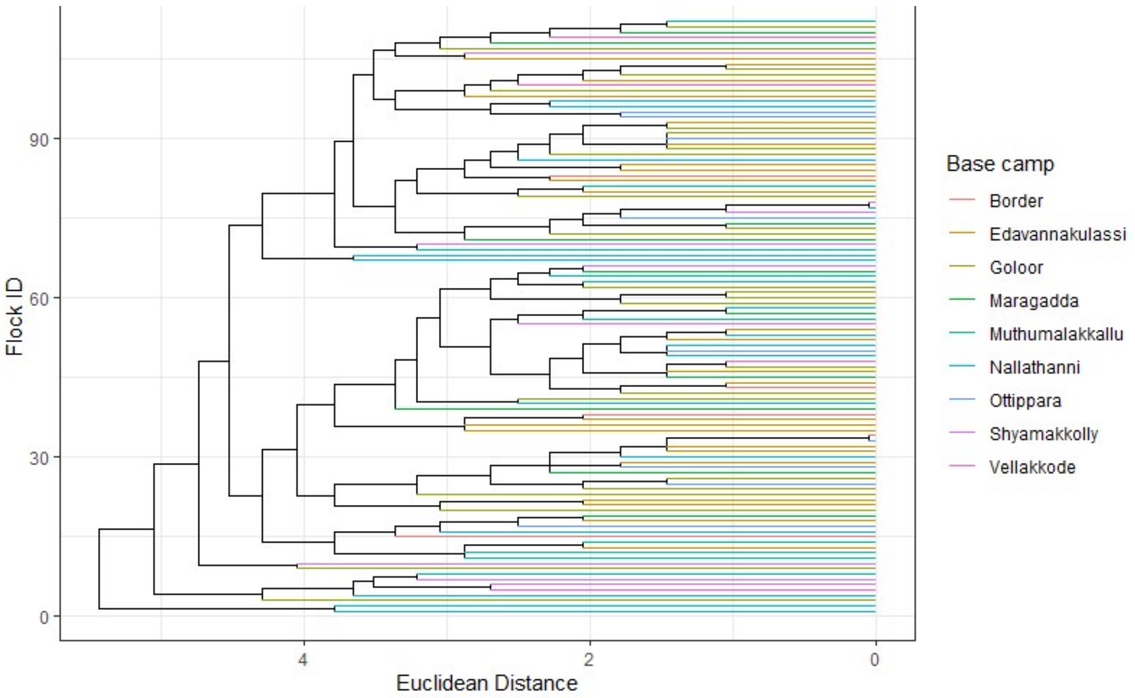
